# Deciphering Hierarchical Chromatin Domains and Preference of Genomic Position Forming Boundaries in Single Mouse Embryonic Stem Cells

**DOI:** 10.1101/2022.05.27.493686

**Authors:** Yusen Ye, Shihua Zhang, Lin Gao, Yuqing Zhu, Jin Zhang

## Abstract

The exploration of single-cell 3D genome maps reveals that chromatin domains are indeed physical structures presenting in single cells and domain boundaries vary from cell to cell. However, exhaustive analysis of regulatory factor binding or elements for preference of the formation of chromatin domains in single cells has not yet emerged. To this end, we first develop a **<u>hi</u>**erarchical **<u>c</u>**hromatin domain **<u>s</u>**tructure identification algorithm (named as **HiCS**) from individual single-cell Hi-C maps, with superior performance in both accuracy and efficiency. The results suggest that in addition to the known CTCF-cohesin complex, Polycomb, TrxG, pluripotent protein families and other multiple factors also contribute to shaping chromatin domain boundaries in single embryonic stem cells. Different cooperation patterns of regulatory factors decipher the preference of genomic position categories forming boundaries. And the most extensive six types of retrotransposons differentially distributed in these genomic position categories with preferential localization.

## Introduction

Genome across a wide range of eukaryotic organisms is efficiently packaged and organized into hierarchical chromatin architecture via ubiquitous architectural features, which is critical to gene regulation and dynamical changes in development and disease [1–3]. These basic features consist of chromatin fibers, which fold into chromatin loops, such as enhancer-promoter interactions and architectural loops mediated by CCCTC-binding factor (CTCF) [4]. These fibers further fold into chromatin domains, referred to as topologically associating domains (TADs) or sub-TADs, which are associated with each other to generate chromosomal compartments. Each chromosome occupies a distinct volume or chromosome territory within the nucleus [4–6]. Genome architecture is an integral part of the chromatin landscape that transcription factors (TFs) must navigate to exert their regulatory roles [7]. Although most loci and chromosomes are characterized by a high degree of order and non-randomness, the precise functional roles and formation mechanism of these features remains obscure [8].

Some TFs, cofactors, and histone modifications that correlate with the chromatin structures have been identified to study features of chromatin organization [9–11]. Particularly, TAD boundaries are enriched with multiple factors including CTCF, cohesin, H3K4me3, H3K36me3, transcription start sites, housekeeping genes, etc, suggesting that CTCF binding, high levels of transcription activity, multiple histone modifications, and other regulatory factors may contribute to the formation of chromatin domains in mammals [8]. Although TADs and their associated regulatory factors have been widely identified in multiple species and are highly conserved and stable across different cell types [6, 12], single-cell 3D genome analysis indicated that they display substantial cell-to-cell variation [13, 14]. Therefore, the bulk analysis only reflects properties of ensemble average structures from millions of cells, which may mask chromatin features appearing in a few cells or a single cell and limit our understanding of chromatin structures.

A recent study reveals that domain structures often adopt globular conformation with strongly physical segregation of neighboring domains, and domain boundaries are preferentially located at CTCF- and cohesin-binding sites with a super-resolution chromatin tracing method [13]. More surprisingly, single-cell domain structures persist even after cohesin degradation [13]. These results suggest other TFs or epigenetic factors may contribute to the formation of chromatin domains. Currently, mouse embryonic stem cells (mESCs) have been served as a specific model cell system to elucidate the mechanisms of 3D genome organization and explore the relationship between chromatin domains and gene regulation. Hundreds of TFs and epigenetic modification profiles have been identified in mESCs [9–11]. The above observations push us to investigate the formation of chromatin domains and their relationship with functional elements in single mESCs systematically.

Here, we first develop a **<u>hi</u>**erarchical **<u>c</u>**hromatin domain **<u>s</u>**tructure identification algorithm (named as **HiCS**) from single-cell Hi-C maps, which shows superior performance in both accuracy and efficiency. We reorganize atlas of ChIP-seq for mESCs, and reveal the patterns of hundreds of regulatory factors are significantly either enriched or absented in domain boundaries of single cells, suggesting that, in addition to known CTCF-cohesin complex, Polycomb, TrxG, pluripotent protein families, different types of histone modifications, and other multiple factors could promote the formation of chromatin domains in single mESCs. To further elaborate on cooperation patterns between different types of regulatory factors, and genomic position categories with differential preference forming boundaries drive by these cooperation patterns, we cluster 13 large genomic position categories (consisting of 29 sub-categories) annotated by 7 different regulatory factor clusters (consisting of 27 sub-clusters). The clear patterns provide a detailed view of the preference of these genomic position categories forming domain boundaries. Furthermore, we discover that these genomic position categories are enriched by different cooperation of retrotransposons with preferential localization. Last but not the least, we find that genomic positions enriched by Alu/B2/B4 retrotransposons have higher preference scores for forming boundaries in G1 and ES phases in comparison with MS and LS/G2 phases, whereas genomic positions enriched by L1/ERVK retrotransposons display opposite tendency. In summary, we reveal that multiple types of regulatory factors interplaying with each other in specific genomic positions could affect focal chromatin interactions, thereby changing interaction density or insulation strength of these regions. This further navigates the preference of genomic position forming boundaries, shape hierarchical chromatin domains, and thus regulate gene expression and cell functions, even cell identity in single embryonic stem cells.

## Results

### Overview of HiCS

The key design of HiCS is to convert the problem of the identification of hierarchical chromatin domains into finding peaks of insulation strength at different genome scales. The domain boundaries usually have higher insulation strength than their neighbors and a relatively large distance from any regions with higher strength (**Fig. 1a**). HiCS calculates two metrics for each bin including the insulation strength *ρ* and the minimum distance between the bin and any other bin with higher strength *δ*, and controls the number of peaks to obtain the hierarchical chromatin domains at different scales by *α* (**Fig. 1b and c**). HiCS is super-fast to identify a chromatin hierarchy that the domain of a higher level embraces the multiple smaller ones of a lower level (**Fig. 2b**). Note that high-level boundaries with high *δ* in local regions may have lower insulation strength than low-level ones, and the boundaries in the same level may have different local insulation strengths, HiCS can automatically detect different level domain boundaries based on local background (**Fig. 2b**).

**Figure 1.**
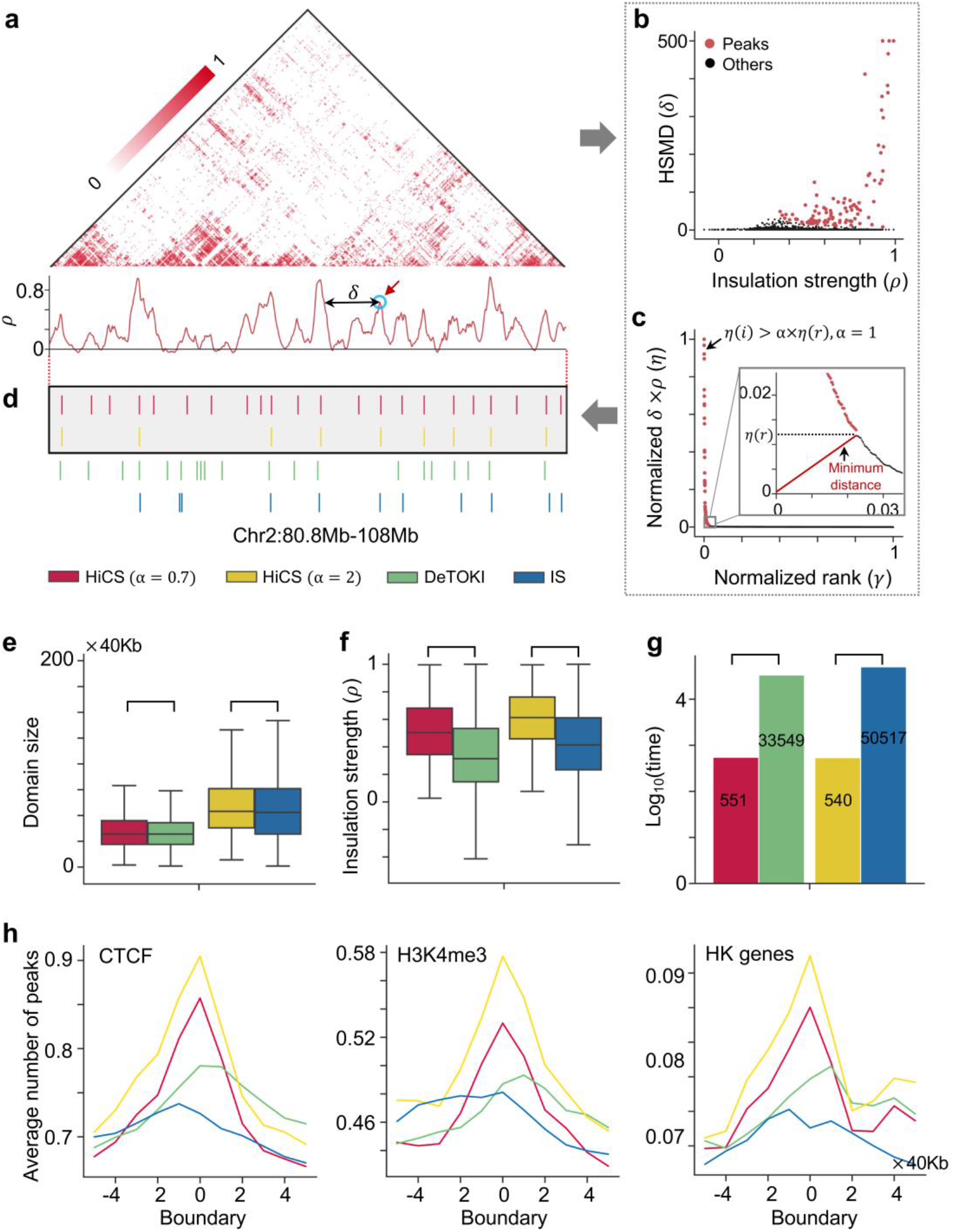
Illustration and efficiency of HiCS for determining the chromatin domains. **a.** An illustrative example of the preprocessed single-cell Hi-C contact map (top) and the insulation strength (*ρ*) of genomic positions (bottom). HSMD (*δ*) represents the minimum distance between the bin and any other bins with higher strengths. **b, c.** The decision graph (b) and the normalization value of *η* = *ρ* × *δ* in a decreasing order (c) for the domain boundaries (colored in red) in the optimal structural identification parameter. The decision graph of optimal structural identification parameter (zoom box in c). **d.** A local example of domain boundaries from map in **a** at the region Chr2:80.8Mb-108Mb with different methods. **e, f.** Comparison of different methods for domain sizes and insulation strengths. **g.** Runtime (seconds) of different methods or parameters. **h.** The average number of CTCF peaks, H3K4me3 peaks, and HK genes at domain boundaries of single cells. The above results (d-h) are implemented by different methods across all single cells.

**Figure 2.**
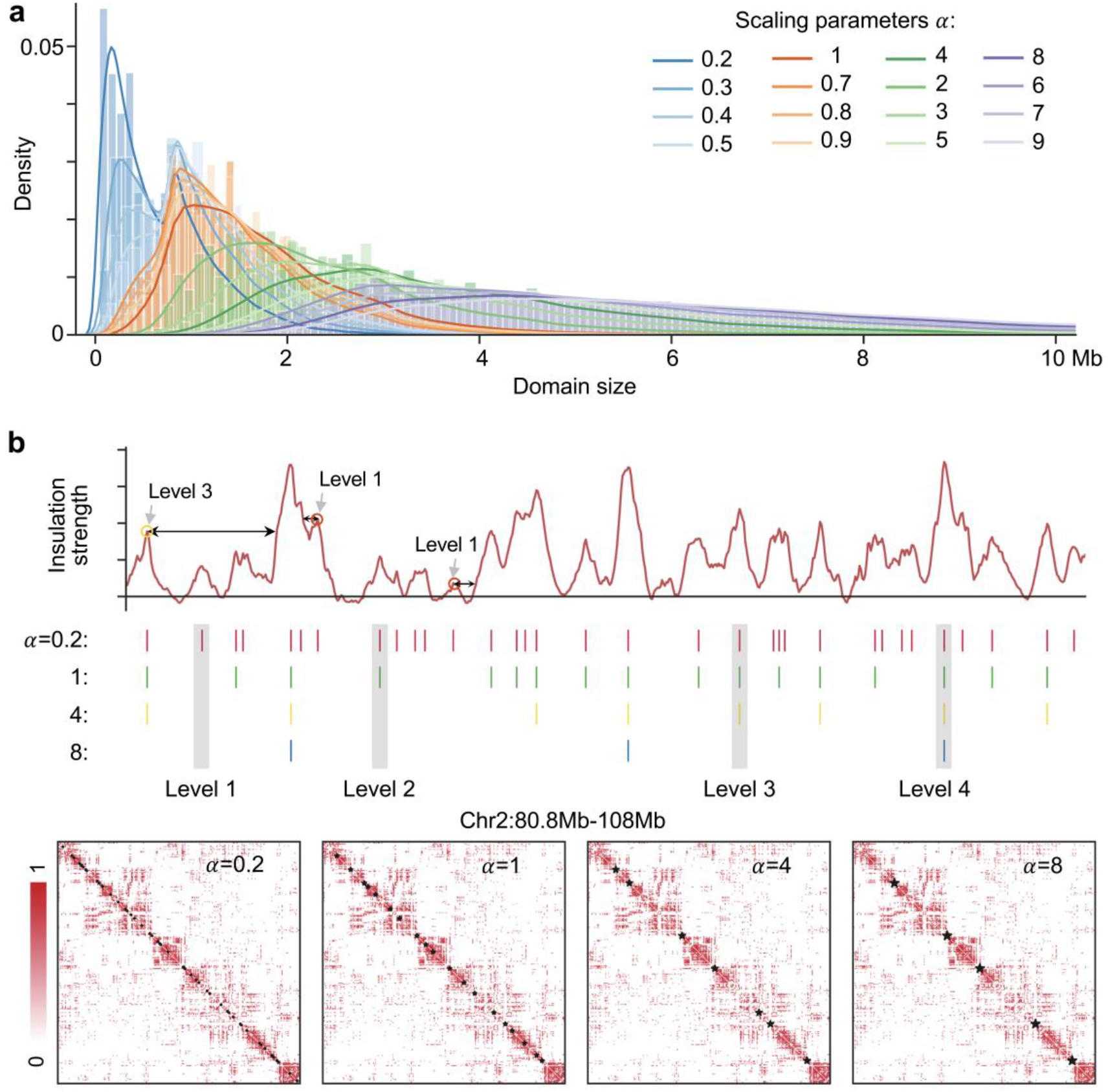
Identification of hierarchical chromatin domains. **a.** Density distribution of domain sizes at different multi-scale parameters. **b.** An example for hierarchical chromatin domains at the region Chr2:80.8Mb-108Mb. Circles mark three boundaries (two boundaries in level 1 have a magnitude difference in local insulation strength, and the boundary in level 3 has lower insulation strength than one of the boundaries in level 1).

Specifically, HiCS consists of three steps (**Methods**): (1) Preprocesses single-cell Hi-C data and calculates the two metrics to find the peaks at a given scale *α*, and determines the domain boundaries for each chromosome of an individual cell (**Fig. 1a-c**); (2) Determines the hierarchical domains of a chromatin by adjusting the optimal structural identification parameter (*α* = 1) (**Fig. 1c**, **Fig. 2a, b**, and **Supplementary Fig. S1a**), Noting that the optimal parameter is automatically determined by the algorithm; (3) Applies a bi-clustering method to group chromatin positions and regulatory factors respectively and analyzes the genomic structure-function relationship by combining hundreds of TFs and epigenetic factors with chromatin domains (**Fig. 4**).

### HiCS shows superior performance

We benchmark the performance of HiCS for domain detection against two methods: one is the commonly used method for bulk data, insulation score (IS) [15], and another is a recent single-cell TAD detection method, deTOKI [16]. We apply these methods to the preprocessed single-cell Hi-C data generated from mESCs [17]. To compare the performance of them fairly, we adjust the scaling parameter of HiCS to obtain similar number and size of domains with IS and deTOKI, respectively (**Fig. 1e**). Actually, the insulation strengths of domains obtained from HiCS is significantly higher than those of IS and deTOKI respectively with similar number of domains, suggesting its superiority to competing methods (**Fig. 1f**). The running time of HiCS is significantly less than both algorithms under the same hardware condition (**Fig. 1g**). Moreover, the domain boundaries detected by HiCS are more significantly enriched in multiple common factors, including CTCF, H3K4me3, Housekeeping (HK) genes (**Fig. 1h**), as well as RNA polymerase II (PolII), promoters, highly expressed genes, and average phastcon score (**Supplementary Fig. S1b**). Also, the boundaries detected by HiCS are more pronounced disappearance of the enhancers or the super enhancers (SEs), which unfavorably form boundaries in single cells as reported on analysis of bulk-cells [9] (**Supplementary Fig. S1b**). With an example, we can see that HiCS can obtain more accurate chromatin domain boundaries at single-cell resolution (**Fig. 1a and d**). Taken together, HiCS shows superior performance in both accuracy and efficiency.

### The existence of hierarchical chromatin domains

We adjust the optimal structural identification parameter to generate multiple-scale chromatin domains at different genomic scales for 1315 single mESCs at 40kb resolution. We clearly observe four peaks of domain size and insulation strength distribution with different scaling parameters of 0.2, 1, 4, and 8, which we choose for the downstream analysis (**Fig. 2a, b** and **Supplementary Fig. S1a**). The domain scales of these four levels are approximately 200Kb~600Kb, 800Kb~1Mb, 2Mb~3Mb, and ~5Mb respectively. We show an example for hierarchical chromatin domains, which well match the local insulation strength of chromatin regions (**Fig. 2b**).

The median size of chromatin domains increases and the median boundaries’ insulation strength enhances with the increase of genomic position level (**Supplementary Fig. S2a-c**). The boundaries show obvious cell-to-cell heterogeneity with a nonzero probability of being located at any genomic positions. 99.8% of genomic positions form boundaries in at least 1% of cells, while only 4.1% of genomic positions form boundaries in more than 14% of cells (**Fig. 3a**). We also observe that the probability forming boundaries enhances with the levels of genomic position increasing (**Supplementary Fig. S2d**). The above observations suggest that domain boundaries vary from cell to cell with nonzero probability at all genomic positions as reported in [13], and the preference of genomic position forming boundaries may shape the formation of hierarchical chromatin domains in single cells.

**Figure 3.**
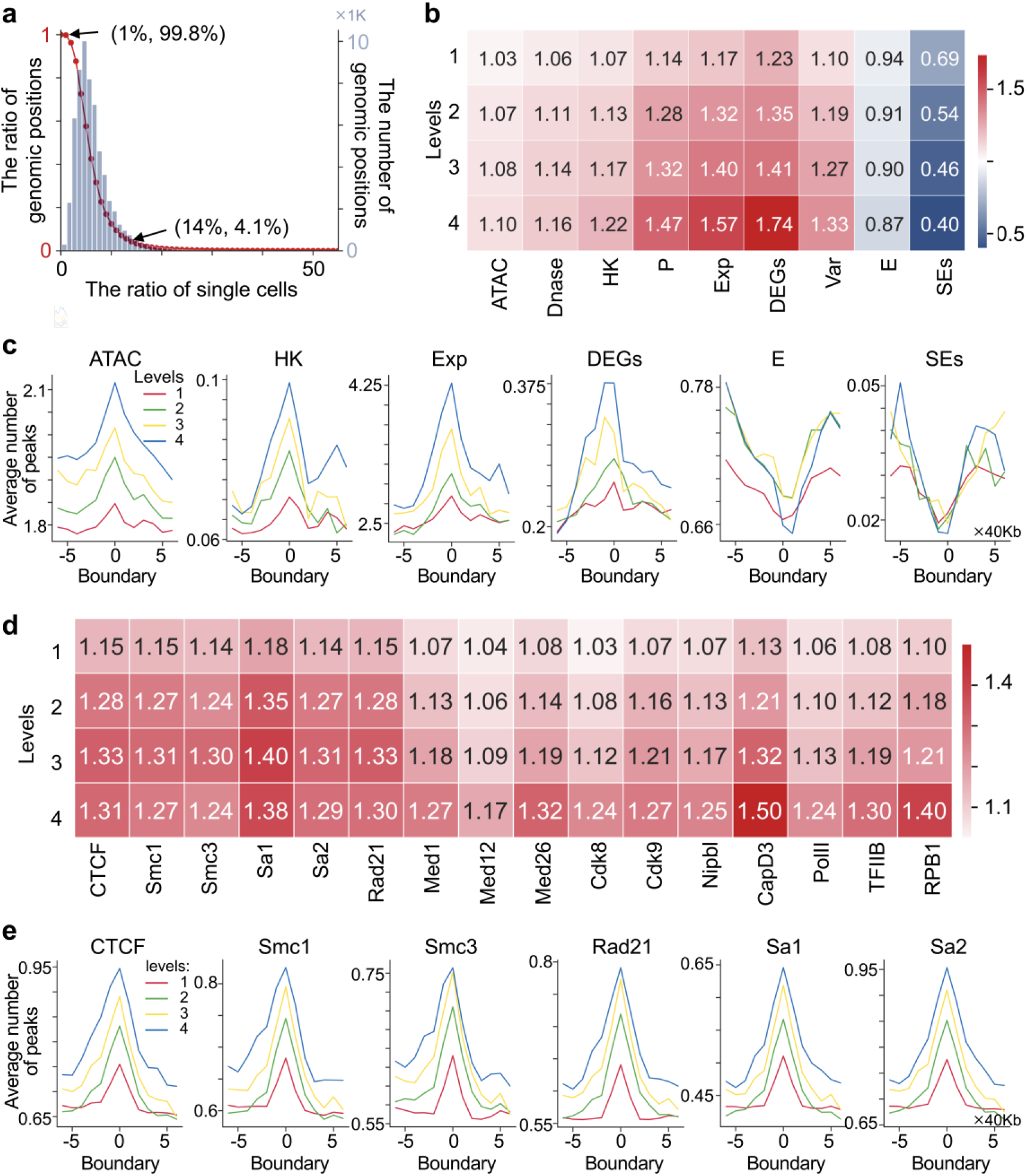
Regulatory factors navigate the preference of genomic position forming boundaries. **a.** The ratio distribution of genomic positions forming boundaries at less than a given ratio of single cells in the left y-axis (such as 99.8% of genomic positions form boundaries in at least 1% of cells, respectively), and the number distribution of genomic positions forming boundaries in the right y-axis as the ratio of single cells increases. **b-c.** The concentration scores (b) and the average number of ATAC-seq peaks and multiple regulatory elements (ATAC: ATAC-seq peaks, HK genes, Exp: gene expression value, DEGs: differentially expressed genes, E: enhancers, SEs) (c) around domain boundaries. **d.** The concentration scores for CTCF, cohesin, mediator- and Pol II-associated factors. **e.** The average number of peaks for CTCF and cohesin around domain boundaries. In b-e, the results were detected in the different genomic scales across all single cells. The concentration score is defined in **Supplementary Methods**.

### Regulatory factors navigate the preference of genomic position forming boundaries

Recent study has shown that domain boundaries are preferentially located at CTCF- and cohesin-binding sites with a super-resolution chromatin tracing method [13]. CTCF and cohesin have been proven to be key factors controlling the functional architecture of mammalian chromosomes forming TADs or sub-TADs by “loop extrusion” [18]. Indeed, we observe that both CTCF and cohesin show similar enrichment patterns in boundaries of single cells in different genomic levels, and as the level of boundaries increases, the degree of enrichment gradually increases (**Fig. 3d and e**).

Previous study has illustrated that high levels of transcription activity may contribute to TAD formation in bulk analysis [19]. Here, we observe that the accessibility of genomic positions, the enrichment degree of HK genes and differentially expressed genes (DEGs), and the expression of genes gradually increases along with the increasing levels of genomic positions (**Fig. 3b, c and Supplementary Fig. S2e**). Conversely, enhancers and super enhancers (SEs) greatly become more absented with the increase of levels (**Fig. 3b and c**). It suggests that the emergence of highly transcriptional activity, especially DEGs, and the absence of enhancers, especially SEs, improve the probability of genomic positions forming boundaries of single cells (**Fig. 3b and c**).

Although the CTCF-cohesin complex is critical for the formation of TADs in mammalian cells, a substantial number of boundaries remain unaffected after cohesin degradation in single cells, suggesting other modulators exist on domain boundaries [7, 20]. We indeed observe that mediators (*Med1*, *Med12*, *Med26*, *Cdk8*, *Cdk9*), *Nipbl*, PolII, TFIIB are all enriched in the boundaries of single cells, and as the level of boundaries increases, the degree of enrichment gradually increases (**Supplementary Fig. S2g-f**). Mediators are essential coactivators that are recruited to the regulatory regions of active genes and facilitate the ability of enhancer-bound TFs to recruit PolII to the promoters of target genes, and *Nipbl* has been proven to bind mediator to load cohesin [9, 10, 21, 22]. The above results suggest that these factors may play important roles in shaping preference of genomic position forming boundaries of single cells.

The master factors of Polycomb repressive complex 1 (PRC1) and PRC2 just exhibit two different enrichment patterns at the domain boundaries of single cells. One type (*Aebp2*, *Rybp*, *Ring1b*) forms obvious single peaks at the domain boundaries, and the degree of enrichment gradually increases with the boundary level increasing (**Supplementary Fig. S3a**). Another type (*Ezh2*, *Pcl2*, *Suz12*, *Eed*) shows double peaks around the domain boundaries (**Supplementary Fig. S3b**). These two types of complexes have been proved to have distinct catalytic activities, but both are generally associated with transcriptional silencing [23]. We also observe that TrxG associated proteins (COMPASS: *Set1a*, *Mll2*, *Mll3/4*, and SWI/SNP: *Brg1*) are all enriched in domain boundaries, except for *Mll3/4* (**Supplementary Fig. S3c**). In mammalian, *Set1a* reportedly contributes to most of the H3K4me3, and *Mll2* mediates H3K4me2 and H3K4me3 at developmental genes, while *Mll3/4* implements monomethylation of H3K4 at enhancers [24]. *Brg1* is an ATP-dependent chromatin remodeler, contributing to the maintenance of pluripotency and self-renewal in ESCs [9]. The above results suggest that the PRC and TrxG protein families could change focal chromatin interactions in different ways.

We find that the core TFs (*Oct4*, *Sox2*, and *Nanog*) controlling the pluripotent state do not prefer to appear in domain boundaries, which may be related to chromatin hubs occupied by super-enhancers/enhancers (**Supplementary Fig. S4a**) as reported in [25]. In addition, we also collect 14 additional TFs that may contribute to the pluripotent state of mESCs and investigate whether they are enriched in domain boundaries of single cells [9] (**Supplementary Fig. S4b and c**). The results indicate that 9 additional TFs (*Esrrb*, *Nr5a2*, *Klf4*, *Zfp281*, *Tcf3*, *Tcfcp2l1, Stat3*, *Prdm14*, and *Smad2/3*) that were previously shown to occupy both typical enhancers and SEs do not prefer to appear in domain boundaries, while 5 factors (*c-Myc*, *n-Myc*, *Zfx*, *Tbx3*, and *Yy*1) that were previously shown to occupy promoter-proximal sites are enriched in domain boundaries [9]. Among them, it is particularly interesting that *Smad2/3*, *Stat3*, and *Tcf3* signaling pathways were considered as key modulators controlling mESCs pluripotent state transition by modifying chromatin states and shaping chromatin domains [26–28].

In addition, we also observe additional 45 proteins are either enriched or absented in domain boundaries of single cells with different enrichment patterns (**Supplementary Figs. S5-7**), which suggests their potential in shaping chromatin domains.

The above TFs and chromatin regulators have the most profound impact on cell states through collaborative control of chromatin states and spatial structures. 14 different histone-modifying enzymes show a variety of enrichment patterns around domain boundaries of single cells (**Supplementary Fig. S8**). For example, ZC3H11A shares consistent enrichment patterns with CTCF-cohesin (**Fig. 3e and Supplementary Fig. S8**). H3K27me3 shares similar enrichment patterns with PRC2, while H2AK119ub1 shares similar enrichment patterns with PCR1 (**Supplementary Figs. S3a and b**), which are consistent with their function in participating chromatin modifications [23]. H3K4me1 marking of enhancers is not enriched in domain boundaries. And H3K4me2 shares similar enrichment patterns with H3K4me3 around domain boundaries, which all are associated with TrxG protein family [24] (**Supplementary Fig. S3c**). It suggests different histone modifications may cooperate with different TFs and chromatin regulators, to modify chromatin states, shape the local chromatin interaction status and organize chromatin domains of single cells.

To sum up, we have observed hundreds of TFs, chromatin regulators, and histone modifications are significantly either enriched or absented in domain boundaries of single cells with differential enrichment patterns. The occupancy of these regulatory factors in specific genomic positions will affect focal chromatin interactions, thereby changing the interaction density or insulation strengths of these regions. These processes may navigate the preference of genomic position forming boundaries, and then shape hierarchical chromosome domains of single cells.

### Cooperation among regulatory factors differentiate genomic position categories

To further elaborate on cooperation patterns between different types of regulatory factors, and genomic position categories with differential preference forming boundaries drive by these cooperation patterns, we grouped 13 large spatially organized genomic position categories (consisting of 29 sub-categories), which were annotated by 7 different regulatory factor clusters (consisting of 27 sub-clusters). The result helps to explain the preference of genomic position forming boundaries in single cells, and providing an increasingly complex view of the genomic structure-function relationship (**Fig. 4 and Supplementary Fig. S9**).

**Figure 4.**
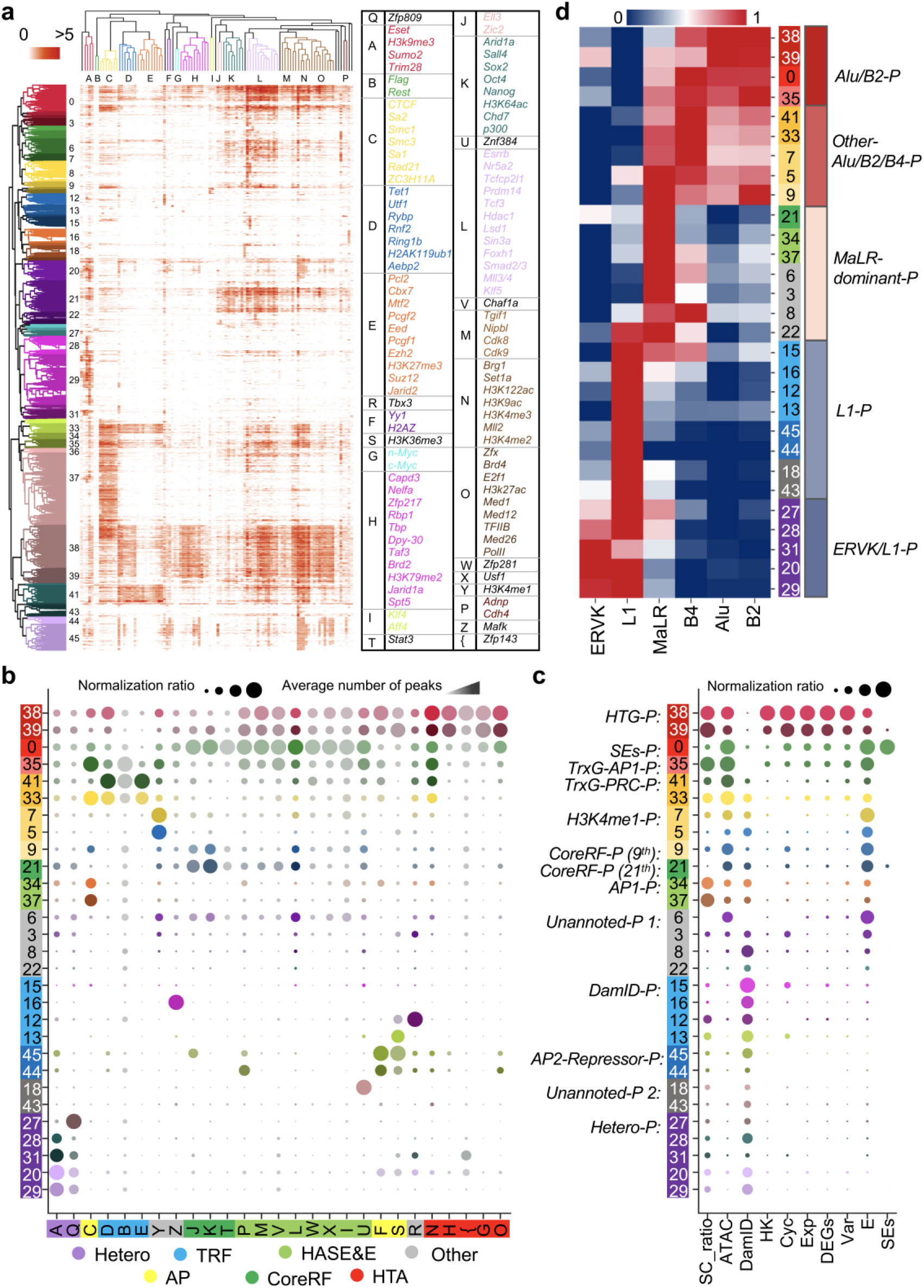
The systematic analysis on genomic position categories. **a.** The bi-cluster of chromatin positions and regulatory factors. **b.** The column normalization ratio and the average number of peaks for regulatory factor-classes on genomic position categories (different dot color represents different categories in Fig. 4a). **c.** The column normalization ratio of different elements or factors, including SC_ratio (the ratio of single cells forming boundaries), ATAC, DamID (DamID-seq), HK genes, Cyc (mark genes for cell cycle), Exp (gene expression value), DEGs, Variable (variable scores for genes), E (Enhancers), and SEs, on genomic position categories. **d.** The heatmap shows the column normalization ratio of different retrotransposons based on genomic position categories.

#### The categories of regulatory factors

The result of hierarchical clustering indicates that these categories mainly consist of the cluster of core regulatory factors of mESC (CoreRF, particularly *Oct4*, *Sox2*, and *Nanog)*, the clusters associated with highly active activators for SEs and enhancers (HASE&E), the clusters of highly transcribed activators (HTA), the clusters of transcriptional repressor factors (TRF), the cluster associated with heterochromatin factors (Hetero), and the clusters of architectural proteins (AP). There are also a few single factors (Other) (**Fig. 4a and Supplementary Fig. S9**). For a specific example with less prior studies among these clusters, *Sumo2* is required to play critical roles in the canonical *Zfp809/Trim28/Eset* complex via post-translational sumoylation of *Trim28*, which enhances the recruitment of *Trim28* to the proviral DNA, resulting in the modification of proviral chromatin with repressive histone H3K9me3 marks in turn [29]. These four factors were grouped with H3K9me3 together in the analysis below, which may organize the formation of heterochromatin (**Fig. 4a**). The annotation information and supporting materials of these all categories were summarized in **Supplementary Table S2**.

We also observe some interesting cooperative patterns among different clusters (**Supplementary Fig. S9b**). For example, the CoreRF cluster is strongly associated with the HASE&E cluster, but not the HTG cluster, while the HASE&E cluster is intimated to the HTG cluster. It suggests the HASE&E cluster may be a bridge between CoreRF and HTG clusters, which can be linked to various signaling pathways involved in the transcriptional network in mESCs.

#### The division of genomic position categories

We further analyze the preference of genomic position categories with a ratio>1% (**Supplementary Fig. S10a**). These genomic position categories can be divided into three families, including the categories with high accessibility preferentially forming boundaries, the categories with high accessibility unfavorably forming boundaries, and the categories with low accessibility unfavorably forming boundaries (**Fig. 5a** and **Supplementary Fig. S10b**).

**Figure 5.**
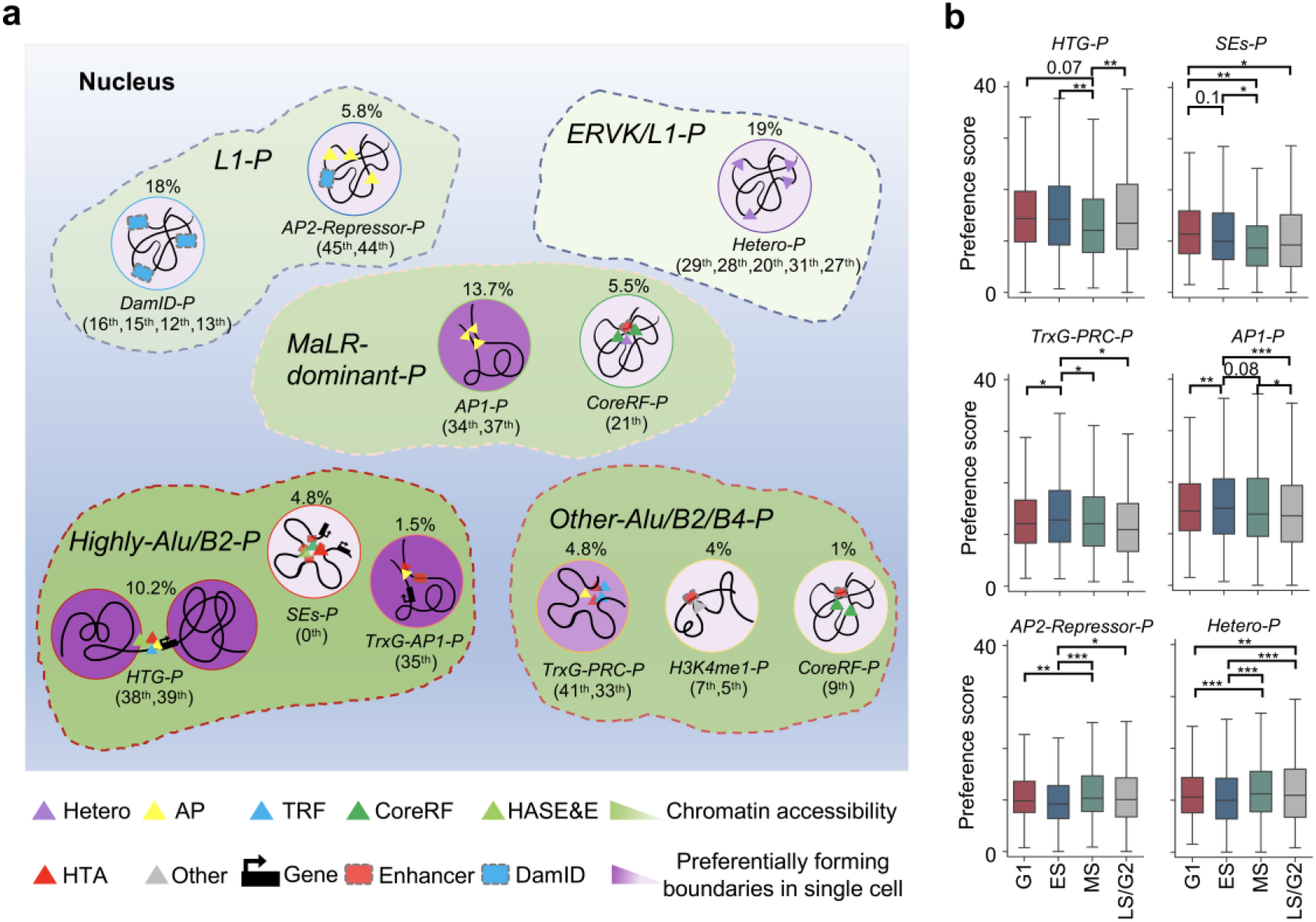
A schematic showing different genomic position categories. **a.** The graph presents the different clusters based on retrotransposons in the area marked by different colors of dotted line containing multiple functional groups associated with different regulatory factor-classes (the sub-categories serial number in above and the ratio occupying from all positions are marked for each functional group). The shades of purple on the background of circles indicate the preference of genomic positions forming boundaries in single cells for each functional group, and the shades of green on the background of these areas marked by dotted line indicate the degree of chromatin accessibility of genomic positions for each cluster based on retrotransposons. **b.** Preference scores of different chromatin landscapes categories across different cell states. Statistical significance is calculated by Welch’s t-test (*P < 0.05, **P < 0.01, and ***P < 0.001). The preference score is defined in **Supplementary Methods**.

Of note, the largest type with weak open chromatin accessibility (13.7% of all positions, consisting of 12.6%, 1.1% from the 37^th^, 34^th^ classes, respectively, defined as AP1-Positions) preferentially forming boundaries only are significantly occupied by AP1 factors (CTCF, cohesin, etc.) comparing with other types of factors (**Fig. 4b, c** and **Fig. 5a**). Previous studies have indicated that loop extrusion executed by CTCF and cohesin is a leading factor governing domain formation and facilitating chromatin folding. We find that the other subunits (*Sa1* and *Sa2*) of cohesin which occupy the same genomic locus and present similar enrichment patterns at boundaries of single cells, suggesting that both *Sa1* and *Sa2* may also participate in the maintenance of domain boundaries (**Fig. 3e**). ZC3H11A, a zinc finger protein, shows a uniform pattern as *Sa1* and *Sa2*, implying its novel role in shaping boundaries of chromatin domains (**Supplementary Fig. S8**). The chromatin positions that AP1 proteins occupy may be directly related to the macroscopic architecture of chromatin within the nucleus, and indirectly change the local chromatin context to exert regulatory functions.

The second major category of open chromatin positions preferentially forming boundaries (7.5%, 2.7% from the 38^th^, 39^th^ classes, defined as HTG-Positions) is associated with highly transcribed genes (**Fig. 4b, c** and **Fig. 5a**). These regions are mainly occupied by the HTA (particularly the TrxG class, playing important roles in orchestrating the stable activation of gene expression) and HASE&E clusters, but not the CoreRF cluster, which indicates that the CoreRF cluster hardly participates in the regulation of genes occupying in domain boundaries of single cells.

The third major category of open chromatin positions preferentially forming boundaries (3.1%, 1.7% from the 41^th^, 33^th^ classes, defined as TrxG-PRC-Positions, 1.5% from the 35^th^ class, defined as TrxG-AP1-Positions) is associated with PRC, TrxG, and AP1 proteins (**Fig. 4b, c** and **Fig. 5a**). The 33^th^ class of genomic regions is also associated with AP1 proteins with a higher probability of forming domain boundaries of single cells compared with the 41^th^ class, which suggests CTCF-cohesin complexes may help these regions form a more stable structure. In addition, the 35^th^ class of genomic region with higher enrichment of TrxG proteins and lower enrichment of PRC proteins compared with the 33^th^ class, which results in higher open chromatin accessibility, enhancers occupation, and probability forming domain boundaries in single cells. The above results indicate that chromatin modifiers (TrxG and PRC proteins) can provide an additional layer of regulation by changing chromatin structures, and balance the formation of domain boundaries by repressing and activating chromatin states, respectively.

The major categories of the open chromatin positions unfavorably forming boundaries are associated with highly active SEs or enhancers, which mainly contain three categories (**Fig. 4b, c** and **Fig. 5a**). The first category (4.8% from the 0^th^ class, defined as SEs-Positions) almost encompasses more than 75% of SEs, which are strongly related with the HASE&E, CoreRF, TrxG, and Med-PolII clusters. The second category (5.5%, 1% from the 21^th^, 9^th^ classes, defined as CoreRF-Positions) is mainly associated with enhancers occupied by CoreRF. The slight difference between the 21^th^ and 9^th^ classes is that the 9^th^ one shows higher chromatin accessibility, while the 21^th^ one shows a stronger DamID enrichment, which may be related to their locations in the nucleus. The third category (2.7%, 1.3% from the 7^th^, 5^th^ classes, H3K4me1-Positions) are related to enhancers enriched by H3K4me1. These regions taking SEs or enhancers associated with CoreRF and H3K4me1 as focal regions generate dense chromatin structures mediated by different regulatory factors in an orientation-independent manner, and unfavorably form boundaries in single cells with weakened insulation strengths [30].

We also check the classes of genomic positions with low chromatin accessibility, all of which unfavorably form boundaries in single cells. These chromatin positions are mainly divided into the following categories (**Fig. 4b, c** and **Fig. 5a**). The first category (19% of all positions, consisting of 9.2%, 3.7%, 3.5%, 1.6%, and 1% from the 29^th^, 28^th^, 20^th^, 31^th^, and 27^th^ classes, respectively, defined as Hetero-Positions) are associated with the Hetero cluster. The second category (18% of all positions, consisting of 2.8%, 2.3%, 1.9%, and 1.7% from the 16^th^, 15^th^, 12^th^, and 13^th^ classes, respectively, defined as DamID-Positions) maintains high enrichment of the DamID signal. The above observations suggest that the presence of heterochromatin factors reduce the probability of single-cell domain boundaries formation. Finally, we observe an interesting and specific category (4.6%, 1.2% from the 45^th^, 44^th^ classes, defined as AP2-Repressor-Positions), which are associated with the AP2 (*Yy1* and H2AZ) and H3K36me3 classes (**Fig. 4a** and **Fig. 5a**). Both *Yy1* and H2AZ facilitate the organization of genome architecture (**Supplementary Table S2**) [31–34].

To summarize, we obtain the following key results by the above analysis: (1) The genomic positions occupied by architectural proteins (CTCF and cohesin), highly transcribed genes, and TrxG proteins preferentially form boundaries in the single cells analysis. (2) The genomic positions taking SEs or enhancers associated with CoreRF or marked by H3K4me1, heterochromatin factors, and repressing factors unfavorably form boundaries in single cells with weakened insulation strengths.

### Retrotransposons are associated with these genomic position categories

In a recent paper, Shen and colleagues find that retrotransposons embedded in 3D genome architecture, regulates the formation of euchromatin and heterochromatin respectively, particularly the separation of compartments A/B [35, 36], which are consistent with our observations. Notably, our study focuses on the effect of retrotransposons on the preference of genomic position forming boundaries in single cells. In order to explain the preference of genomic position categories forming boundaries in single cells with more detail, we further analyze the influences of the largest six types of retrotransposons on genomic architecture.

We observe that the division of genomic position categories is strongly associated with the regulation of retrotransposons. The genomic positions are clearly divided into five functional units based on categories of retrotransposons, including Highly-Alu/B2-Positions, Other-Alu/B2/B4-Positions, MaLR-dominant-Positions, L1-Positions, and ERVK/L1-Positions (**Fig. 4d** and **Fig. 5a**).

Firstly, all genomic regions (Hetero-Positions, DamID-Positions, AP2-Repressor-Positions) with low accessibility and unfavorably forming boundaries are enriched by L1 elements, which tend to occupy gene-poor, heterochromatic B compartments that interact with lamina-associated domains in previous studies [35]. Among the defined regions, we are surprised to find that Hetero-Positions are specially associated with ERVK elements, which may indicate that ERVK acts as specific roles in regulating embryonic development as reported in [37, 38]. The above observation also implies that the previously unannotated 18^th^ and 43^th^ genomic positions may be related to heterochromatin organization.

We also observe that genomic regions (HTG-Positions, SEs-Positions, TrxG-AP1-Positions, TrxG-PRC-Positions, H3K4me1-Positions, the 9^th^ class of CoreRF-Positions) are enriched by Alu/B2/B4, which may be related to euchromatin organization [35]. HTG-Positions, SEs-Positions as well as TrxG-AP1-Positions with higher Alu/B2 enrichment than others suggest that the enrichment of Alu/B2 may indicate the transcription level of genes, and promote the formation of hierarchical chromatin structures by regulating gene transcription and SEs/Enhancer activation.

Besides the above genomic positions, what is interesting is that MaLR elements are enriched in AP1-Positions, the 21^th^ class of CoreRF-Positions, and other unannotated regions (the 6^th^, 3^th^, 8^th^, and 22^th^ classes). We first observe that the 21^th^ class occurs a stronger heterochromatin factors (the Hetero cluster) enrichment compared to the 9^th^ (CoreRF-Positions), which may result in uncertain chromatin states in the 21^th^ positions (**Fig. 4b and c**). In addition, previous studies have shown that domain boundaries mediated by AP1 proteins (e.g., CTCF and cohesin) may block the spread of chromatin states [4]. It may suggest that the regions dominantly enriched by MaLR elements may often undergo switches between euchromatin and heterochromatin.

In summary we find that: (1) L1-Positions with low accessibility and unfavorably forming boundaries are associated with heterochromatin organization, and Alu/B2/B4-Positions are associated with euchromatin chromatin, which are consistent with a recent study [35], while MaLR-Positions may result in switches between euchromatin and heterochromatin, which is yet to be proven. (2) ERVK elements acts as specific roles heterochromatin formation, while Alu/B2 may promote highly transcription of genes and highly activation of SEs/Enhancers. These retrotransposons contribute to the maintaining of chromatin states, and interplay with other types of regulatory factors, to navigate the preference of genomic positions forming boundaries and gene regulation in single cells.

### Genomic landscape regulates cellular states

To investigate the preference of genomic positions in the above functional groups along the cell cycle process, we check the dynamics of each functional group forming boundaries in single cells among four different cycle phases, including G1, early-S (ES), mid-S (MS), and late-S/G2 (LS/G2). We first reveal that genomic positions enriched by SEs show a significant preference for forming boundaries in the G1 phase (**Fig. 5b**). The observation suggests that highly activation of SEs in the phase may promote gene regulation and transcription for cell growth in size, and ensure biomaterials for DNA synthesis. In addition, the functional group (HTG-Positions) accompanied with highly transcribed genes exhibits a significant loss of boundaries in the MS phase, which may be because the rates of transcription and protein synthesis are low during DNA replication (**Fig. 5b**). We also observe that functional groups occupied by both CTCF-cohesin and TrxG-PRC complexes prefer to form boundaries in ES phases (**Fig. 5b**). The observation implicates in the clearest segmentation of chromatin structures at the beginning of DNA replication [39, 40]. Both complexes have been proven to modify local chromatin structure and regulate higher-order chromatin organization [7, 41]. And functional groups (Hetero-Positions and AP2-Repressor-Positions) associated with heterochromatin organization prefers to form boundaries in both MS and LS/G2 phases (**Fig. 5b**). Both phases may prepare for everything entering the mitosis phase with condensing chromatin states.

In general, genomic positions (HTG-Positions, SEs-Positions, TrxG-AP1-Positions, TrxG-PRC-Positions, H3K4me1-Positions, and the 9^th^ class of CoreRF-Positions) enriched by Alu/B2/B4 retrotransposons have higher preference scores for forming boundaries of single cells in G1 and ES phases in comparison with MS and LS/G2 phases, whereas genomic positions (Hetero-Positions, AP2-Repressor-Positions, and DamID-Positions) enriched by L1/ERVK retrotransposons display opposite tendency, following by high preference scores for forming boundaries in both MS and LS/G2 phases (**Fig. 5b** and **Supplementary Fig. S11**). The above observations further expound that the dynamic interplay among different types of regulatory factors, retrotransposons, and chromatin structures could navigate gene regulation and cell functions, even cell identity in single embryonic stem cells.

## Discussion

Several decades of research have shown that eukaryotic chromatin adopts a complex hierarchical architecture within the nucleus, which plays a key role in functional implications for almost all nuclear processes. Thus, the spatially organized chromatin architecture interplaying with multiple types of regulatory factors shape focal chromatin landscapes and then exert gene regulatory functions. Single-cell 3D genome analysis extends the limitation of bulk analysis to show substantial cell-to-cell variation and promote our understanding of chromatin structures in the individual cell. Recent discoveries on single-cell 3D genome have shed light on the relationship between CTCF-cohesin complexes and domain formation, but the more molecular details associated with regulatory factors remain to be investigated [13].

Here, we develop HiCS to detect hierarchical chromatin domains from single-cell Hi-C maps, and observe hundreds of regulatory factors, including TFs, chromatin regulators, and histone modifications, are significantly either enriched or absented in domain boundaries of single cells, which presents several different enrichment patterns. The results suggest their potential cooperative associations in shaping focal chromatin interactions, thereby changing interaction density or insulation strength of these regions, and drive different genomic position categories. We further group chromatin position categories and different regulatory factor clusters, explaining the emergence and functionality of different chromatin landscapes and providing a comprehensive view of the genomic structure-function relationship. We also find that different retrotransposons exactly match the above genomic position categories. The above results indicate that these regulatory factors interplaying with each other exert gene regulatory processes and control cell functions, even cell identity.

The chromatin structures within the nucleus operate in an obvious dynamic process driven by both “loop extrusion” and attractive process induced by regulatory factors (associated with compartmentalization). The process may condense or loose local chromatin landscapes in an overlapping and concerted manner accompanied by adjusting insulation strength of chromatin position and generating chromatin loops, and then shape gene expression programs during cell-fate specification [39]. Further work is needed to leverage more specific chromatin structures, particularly chromatin loops, of single cells with more abundant regulatory factors (TFs, chromatin regulators, histone modifications, retrotransposons, RNA, even structural variations) to understand structure-function relationships in complex tissues or diseases, particularly cancers [42, 43]. It will promote our understanding of how multiple types of regulatory factors interact with chromatin topological engines (such as loop extraction and compartmentalization) to regulate the gene-repression program, determine cell functions and identity, and further explain tissue complexity and disease development.

## Materials and Methods

### Single-cell Hi-C and other genomic data processing

#### Single-cell Hi-C data generated from mESCs

The single-cell Hi-C dataset used in this study consists of 1992 diploid cells of mESCs grown in 2i media without feeder cells with stringent quality control filter. This dataset involves a median number of 393506 restriction fragments, and 127233 distinct >1 kb contacting pairs on average per cell [17]. The top 1315 cells with >250000 contacts per cell were selected for downstream analysis. Among them, 317, 341, 303, and 354 cells belong to G1, early-S (ES), mid-S (MS), and late-S/G2 (LS/G2) phases labeled by fluorescence-activated cell sorting (FACS) sort criterion, respectively.

#### An atlas of ChIP-seq for mESCs

We organized an atlas of ChIP-seq for hundreds of regulatory factors of mESCs (**Supplementary Table S1**), including CTCF, cohesin (*Smc1, Smc3, Rad21, Sa1, Sa2*), mediators (*Med1, Med2, Med26, Cdk8, Cdk9*), codensin (*Capd3, Nipbl*), PolII, TFIIB, polycomb repressive complex (*Aebp2, Rybp, Ring1b, Rnf2, Suz12, Ezh2, Eed, Pcl2*), Trithorax protein family (*Set1a, Mll2, Mll3/4, Brg1*), the core regulatory factors of mESC (*Oct4, Sox2, Nanog*), the regulatory factors of ESC occupied on enhancers or SEs (*Esrrb, Nr5a2, Klf4, Stat3, Prdm14, Zfp281, Tcf3, Tcfcp2l1, Smad2/3*), the regulatory factors of ESC occupied on promoter-proximal sites or sites that border topological domains (*c-Myc, n-Myc, Zfx, Tbx3, Yy1*), and additional 45 TFs. We collected 14 histone modification factors, including H3K4me1, H3K4me2, H3K4me3, H3K27me3, H3K36me3, H3K79me2, H3k9me3, H3K9ac, H3K122ac, H3K64ac, H3K27ac, H2AZ, ZC3H11A, and H2AK119ub1. In addition, we also collected two chromatin accessibility datasets (ATAC-seq and Dnase-seq) and the DamID-seq dataset.

For the ChIP-seq of regulatory factors, peaks were called using MACS2 software with q-value cut-off 1 × 10^−5^ [44]. The source information and supporting materials of these factors were summarized in **Supplementary Table S1 and S2**.

#### Regulatory elements and genes

Enhancers/SEs and gene expression datasets of mESCs were downloaded in GSE29278 [45]. Housekeeping genes were downloaded in Housekeeping and Reference Transcript Atlas (HRT Atlas v1.0, www.housekeeping.unicamp.br) [46]. PhastCons scores were downloaded from the UCSC Genome Browser via ftp://hgdownload.cse.ucsc.edu/goldenPath/mm9/phastCons30way/vertebrate [47]. Mouse cell-cycle annotated genes were obtained from the mouse genome informatics (MGI) (http://www.informatics.jax.org/), containing 891 genes relating to cell cycle process and regulation. We adopted Seurat to detect DEGs and the variable score of genes using *FindAllMarkers* and *FindVariableFeatures* functions based on single-cell RNA-seq data of ESCs, which consists of 182 cells labeled by FACS sort criterion, including 59, 58, and 65 cells belonging to “G1” phase, “S” phase, and “G2M” phase, respectively [48, 49].

#### Retrotransposons

Retrotransposons built from RepeatMasker annotations were downloaded from the UCSC Table Browser (http://genome.ucsc.edu/). We kept the top six categories of counts for the downstream analysis, including Alu, B2, B4, MaLR, L1, and ERVK.

### HiCS

#### Preprocess contact probability for each chromosome of the individual cell

We first divided each chromosome into bins of specific size (40kb in this study), and counted the contact for each bin pair. Next, we modeled each chromosome as an unweighted network (each bin is one node, and each bin pair with non-zero contacts is added as one edge), and implemented a classic graph embedding method node2vec, which applies a biased random walks procedure, to compute the contact probability of edges by computing the cosine similarity of any two node embedding vectors, and obtained the preprocessed matrix *A* (**Fig. 1a**) [50]. We only kept the top 5% pairs for downstream analysis and the diagonal pairs were removed in our study.

#### Detect domain boundaries for each individual cell

Inspired by a fast density-based clustering method designed for grouping data points [51, 52], we take advantage of finding the cluster centers to detect domain boundaries for each chromosome of individual cells (**Fig. 1b and c**). Specifically, we define two indexes for each 40kb bin: (1) insulation strength *ρ*(*i*) of the i^th^ genomic position is defined as the ratio of (*I*_*i*,*intra*_ − *I*_*i*,*inter*_) and (*I*_*i*,*intra*_ + *I*_*i*,*inter*_) [53] using a 800kb sliding window size:

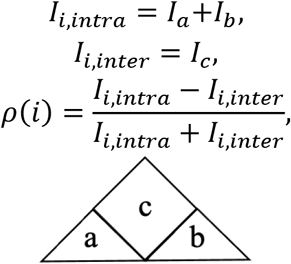

where *I*_*a*_, *I*_*b*_, and *I*_*c*_ respectively represent the summation of interaction frequencies for region *a*, *b*, and c, and (2) minimum distance between the bin *i* and any other bin *j* with higher insulation strength is defined as *δ*(*i*):

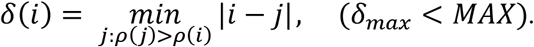

We search for higher insulation strength of bin *i* in the range of *MAX* = 500 (20M genomic distance at 40kb resolution). Next, we define *ρ*^*′*^(*i*) = *ρ*^*′*^(*i*)/*ρ*_*max*_ and *δ*^*′*^(*i*) = *δ*(*i*)/*δ_max_* such that both *ρ^′^*(*i*) and *δ^′^*(*i*) are within range [0,1]. Then, we generate the rank *γ* of all bins for each chromosome by their *η*(*i*) = *ρ*^*′*^(*i*) × *δ*^*′*^(*i*) in the descending and normalized the rank of each bin by *γ*^*′*^(*i*) = *γ*(*i*)/*γ*_*max*_. We define the optimal reflection point with 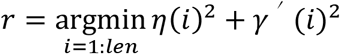, where *len* is defined as the number of bins for a specific chromosome. The boundaries of the optimal structure are assigned by bins with *η*(*i*) > α × *η*(*r*), (*α* = 1). The gap regions are defined by *I*_*i*,*inter*_ = 0 or have no contact with any other bins.

#### Determine the hierarchical chromatin domains

We ran the above procedures of detecting domain boundaries multiple times by define *η*(*i*) > α × *η*(*r*) as a screening selection for different genomic levels, where the range of α are set as (0.1, 10) (**Fig. 2a**). In our study, we selected multiple scaling parameters α as {0.2, 1, 4, 8} to obtain the hierarchical domains of chromatin, according to the distribution of the domain sizes and insulation strengths under different scaling parameters (**Fig. 2a and b**).

#### Clustering and annotating genomic positions with different types of regulatory factors

We applied the hierarchical clustering method to group regulatory factors and chromatin positions, respectively (**Fig. 4a**). The classes of chromatin positions with the ratio>1% of all chromatin positions were selected for downstream analysis, which led to 29 classes of chromatin positions, along with 27 different transcriptional factor classes.

We further applied the hierarchical clustering method to merge the 27 classes into 7 large clusters, based on the Pearson correlation of the normalization ratio of the mean counts for peaks of regulatory factor classes (**Supplementary Fig. S9b**). These clusters or sub-clusters were manually annotated based on the annotation information (**Supplementary Table S2**). We then merged different chromatin position classes into 12 large categories based on hierarchical clustering of correlation of regulatory factor classes, and manually annotated these categories based on the ratios of regulatory factor classes. We obtained and annotated 12 large chromatin position categories (consisting of 29 sub-categories), and 7 different regulatory factor clusters (consisting of 27 sub-clusters).

## Supporting information

Suuplementary Materials

## Data availability

All datasets analyzed in this study were published previously. The corresponding descriptions and preprocessing steps can be found in Supplementary Materials.

## Software availability

The open-source HiCS python package and tutorial are available at GitHub (https://github.com/YusenYe/HiCS).

## Acknowledgements

This work was supported by the National Natural Science Foundation of China [No. 62002275 to Y.Y., Nos. 62132015 & 61873198 to L.G., and Nos. 12126605&61621003 to S.Z.], the National Key Research and Development Program of China [No. 2019YFA0709501 to S.Z.], the Strategic Priority Research Program of the Chinese Academy of Sciences (CAS) [Nos. XDA16021400, XDPB17 to S.Z.].

## Author Contributions

Y.Y. conceived the idea, implemented the algorithm and performed the analyses. Y.Y. interpreted the results. S.Z. and L.G. provided scientific insights on the applications. Y.Y. wrote the manuscript with feedback from all other authors. All of the authors read and approved the final manuscript.

## Completing interests

The authors declare no completing interests.

