## Supplementary material for "Deciphering Hierarchical Chromatin Domains and Preference of Genomic Position Forming Boundaries in Single Mouse Embryonic Stem Cells": Suuplementary Materials

Science Advances
AAAS

Supplementary Materials for

### Deciphering Hierarchical Chromatin Domains and Preference of Genomic Position Forming Boundaries in Single Mouse Embryonic Stem Cells

Yusen Ye[1*], Shihua Zhang[2,3,4], Lin Gao[1], Yuqing Zhu[5], Jin Zhang[6,7,8,9]

**This PDF file includes:**

Supplementary Text
Figs. S1 to S11
Tables S1 to S2

**Supplementary Text**

The columns normalization ratio of different regulatory factors

The columns normalization ratio of the $k^{th}$ factor on the $l^{th}$ chromatin landscape class is defined as

$$CNR(k,l) = \frac{MC(k,l) - \min_{i=1,\dots,m} MC(k,i)}{\max_{i=1,\dots,m} MC(k,i) - \min_{i=1,\dots,m} MC(k,i)}$$

where $MC(k,i)$ represents the average number for peaks of the $k^{th}$ factor on the $i^{th}$ chromatin landscape class.

The concentration scores

The concentration scores of the ith regulatory factor class on domain boundaries across all single cells are defined as

$$CS(i) = \begin{cases} m(b=0)/\min_{k=-6,\dots,6} m(b=k), & if\ m(b=0) \geq m(-l \leq b \leq l) \\ m(b=0)/\max_{k=-l,\dots,l} m(b=k), & else \end{cases},$$

where $m(b=0)$ represents the average number of peaks on domain boundaries across all single cells and $m(-l \leq b \leq l)$ represents the average number of peaks on all $\pm 6$ bins around domain boundaries across all single cells. $m(b=l)$ represents the average number of peaks on the $|k|^{th}$ bin of upstream or downstream of domain boundaries across all single cells.

The preference score of genomic positions across different cell states

The preference score of genomic position $k$ is defined as $PS(k) = c_s^k * 10^6 / \sum_{i=1}^{n} c_s^i$, where $c_s^i$ represents the counts of genomic position $i$ for forming boundaries given one cell state $s$, and $n$ is the number of genomic positions.

#### Supplementary Figures

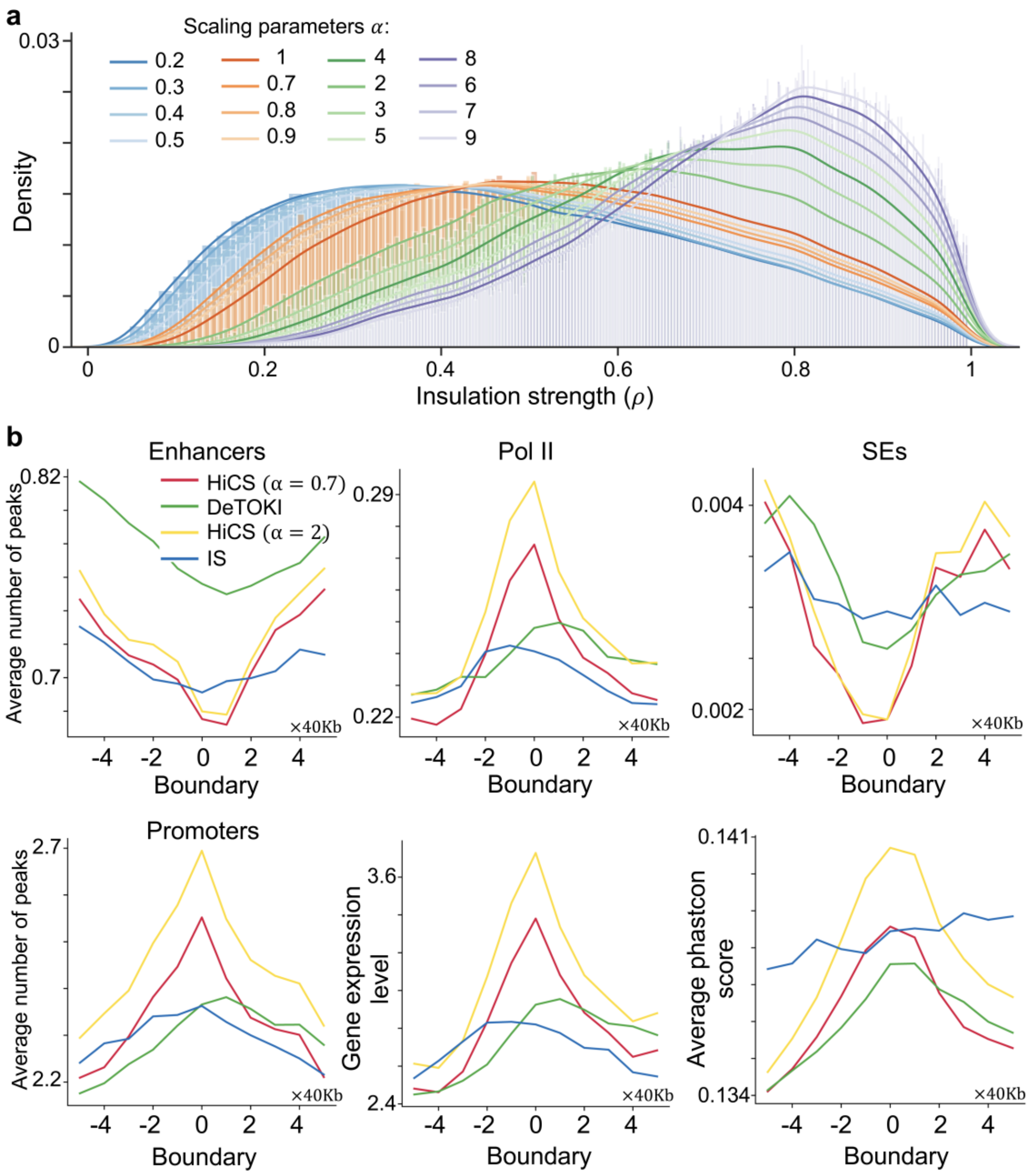


**Fig. S1. a.** The density distribution of insulation strengths at multiple scaling parameters. **b.** The average number of PolII peaks, enhancers, SEs, promoters, the gene expression level, and the mean phastcon score at domain boundaries of single cells. The above results were implemented by different methods or scaling parameters across all single cells.

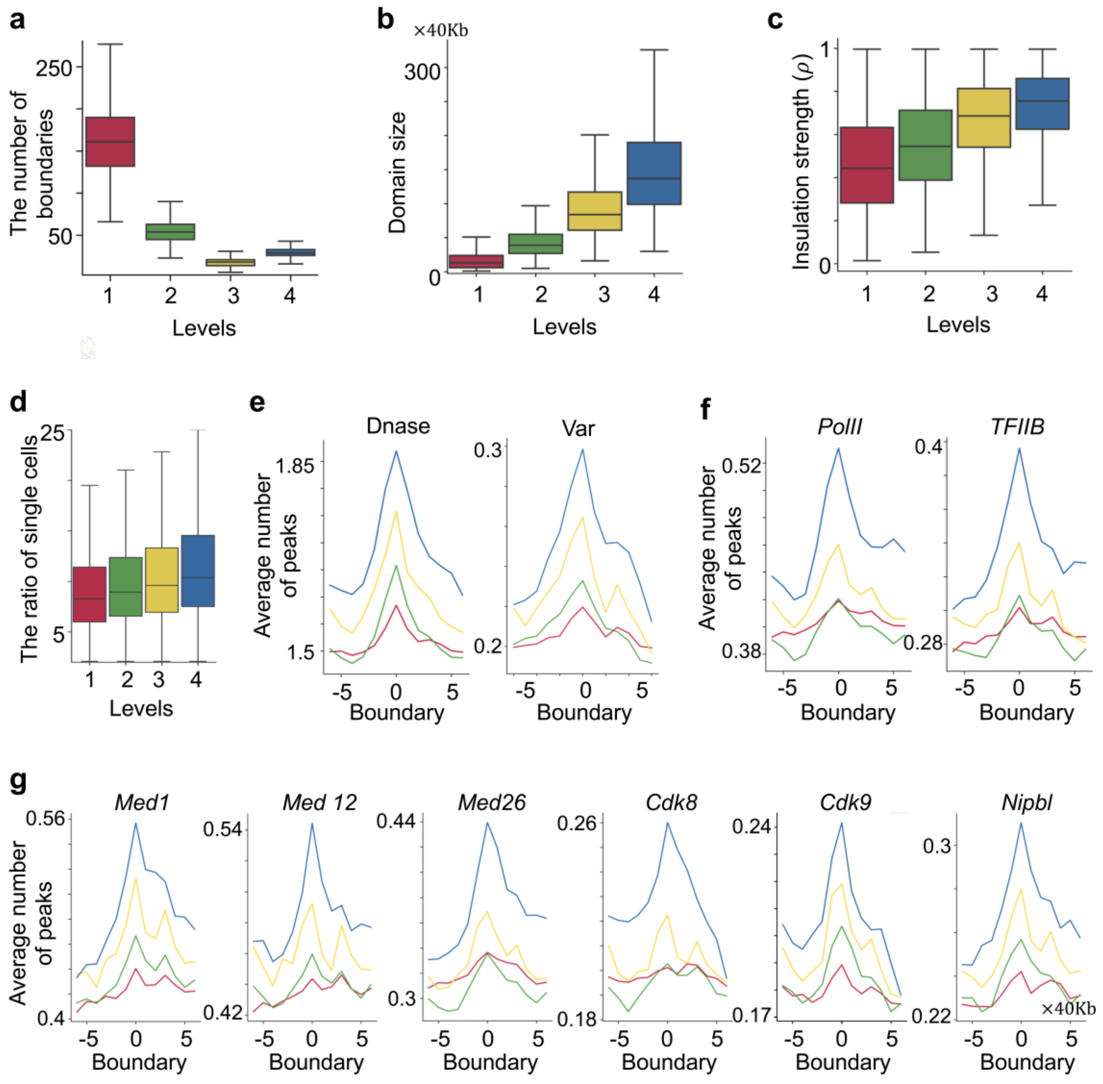


**Fig. S2. a.** The average number of boundaries. **b.** Domain size. **c.** Insulation strengths of boundaries. **d.** The ratio of single cells forming boundaries. In **a-d**, the results were detected in the different genomic scales across all single cells on Chromosome 2. **e-g.** The average number of Dnase-seq peaks, Var (variable scores for genes), and multiple regulatory factors (PolII- and mediator-associated factors) peaks across domain boundaries of all single cells in the different genomic scales.

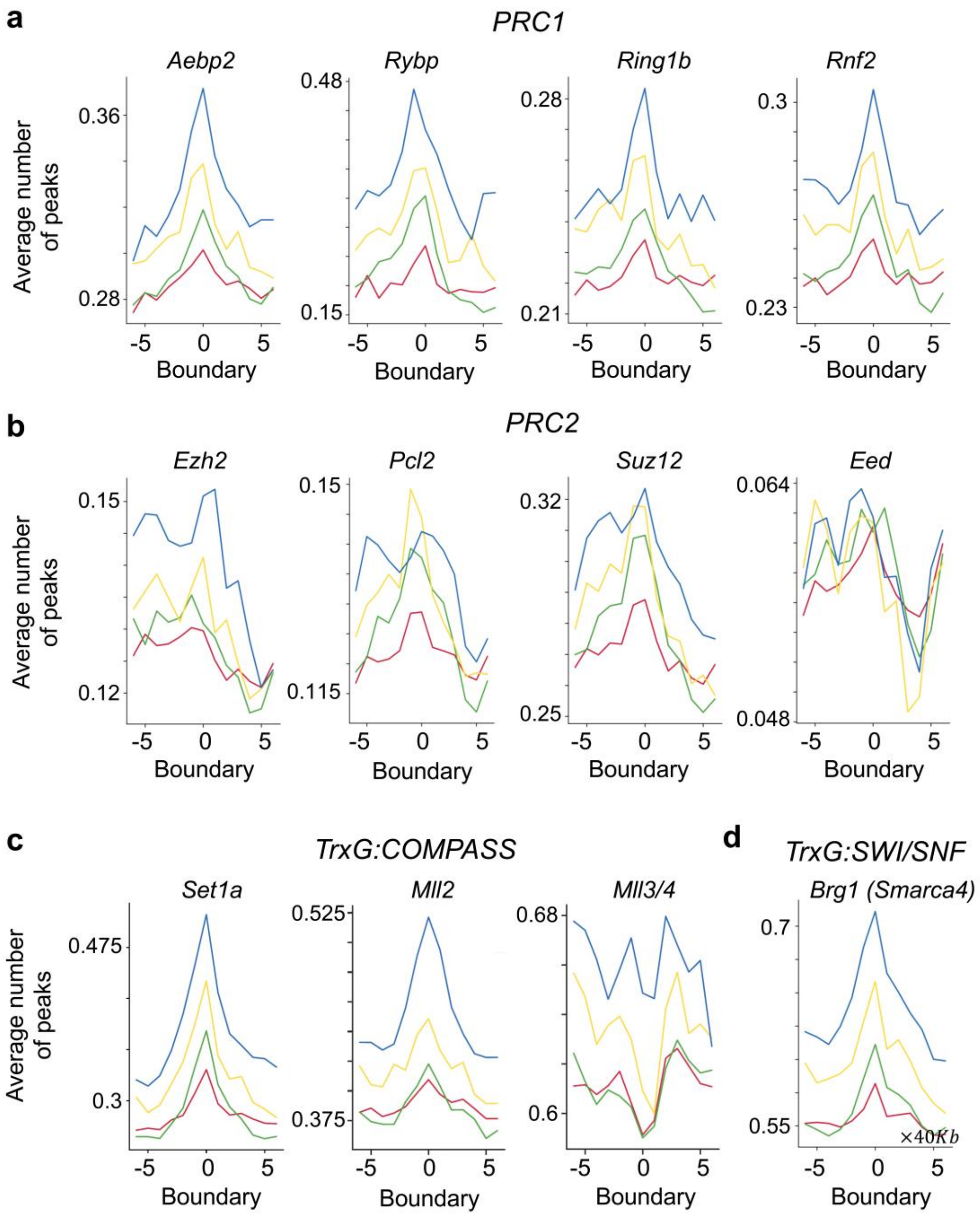


**Fig. S3. a, b, c.** The average number of peaks for the master Polycomb repressive complex 1 and 2(PRC1 (a) and PRC2 (b), respectively) and TrxG proteins (COMPASS and SWI/SNF) (c) across domain boundaries of all single cells in the different genomic scales.

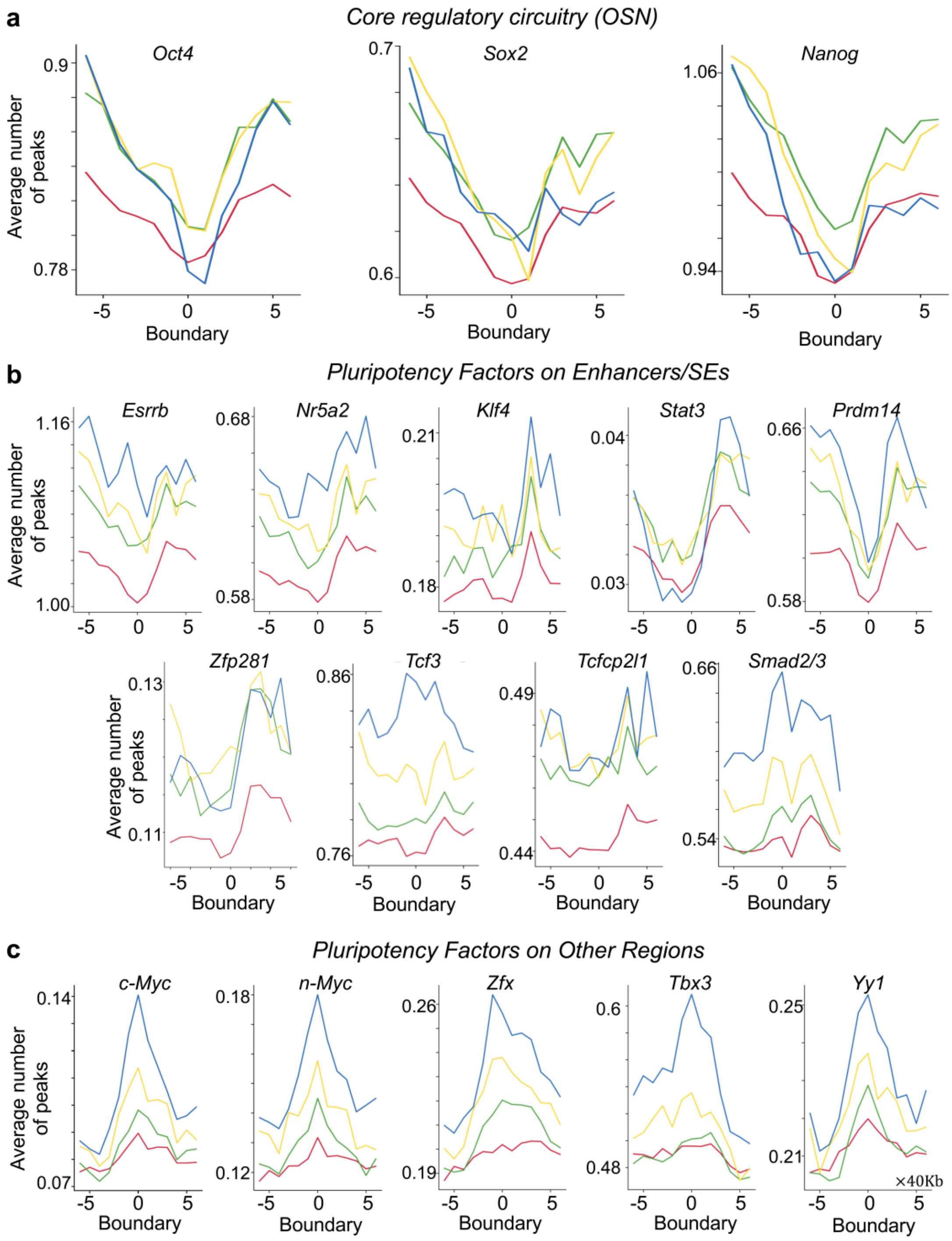


**Fig. S4. a, b, c.** The average number of peaks for the core regulatory circuitry factors (**a**), pluripotency factors enriched in enhancers/SEs (**b**), and pluripotency factors enriched in other regions such as promoter-proximal sites (**c**) across domain boundaries of all single cells in the different genomic scales.

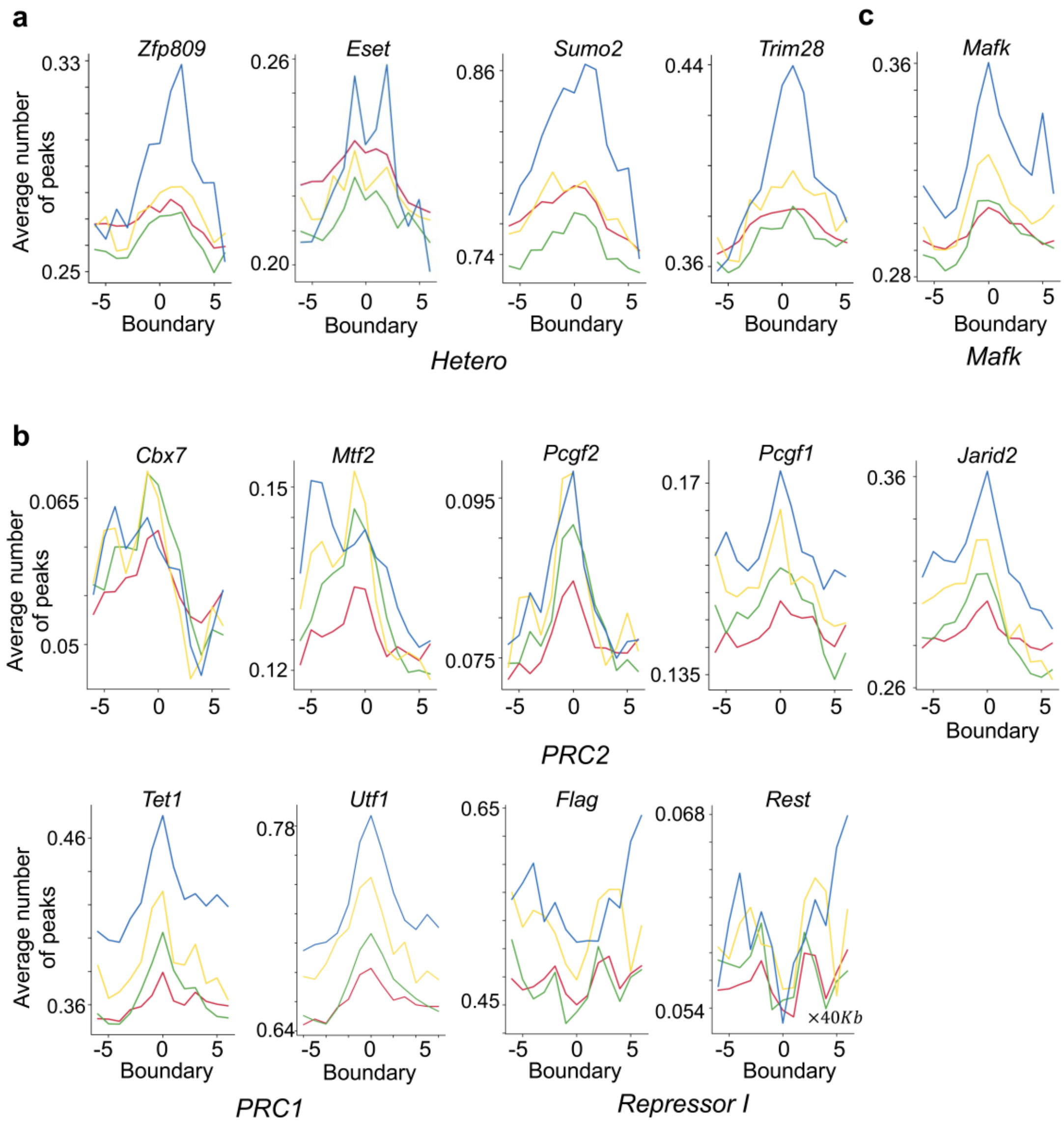


**Fig. S5.** The average number of peaks for multiple histone modifications across domain boundaries of all single cells at the different genomic scales. Each histone modifications correspond to different regulatory factors classes annotated in Fig. 4b.

#### Supplementary Figure S6

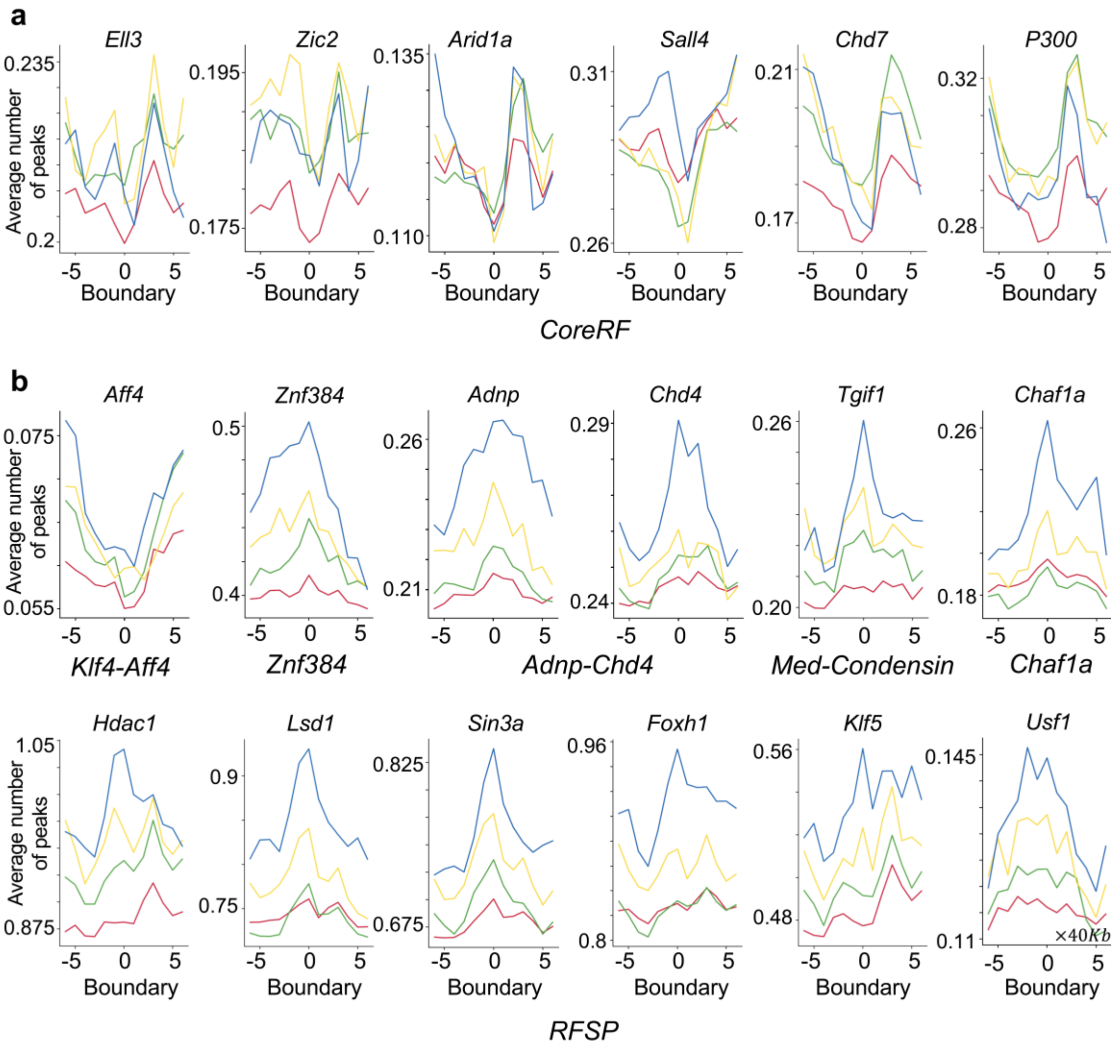

### Supplementary Figure S7

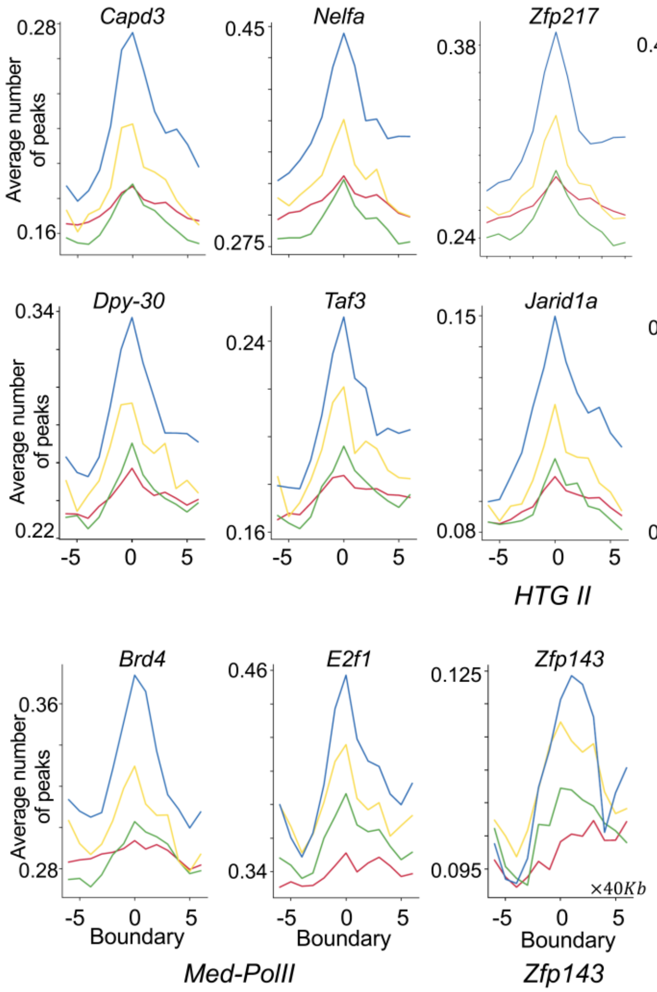

#### Supplementary Figure S8

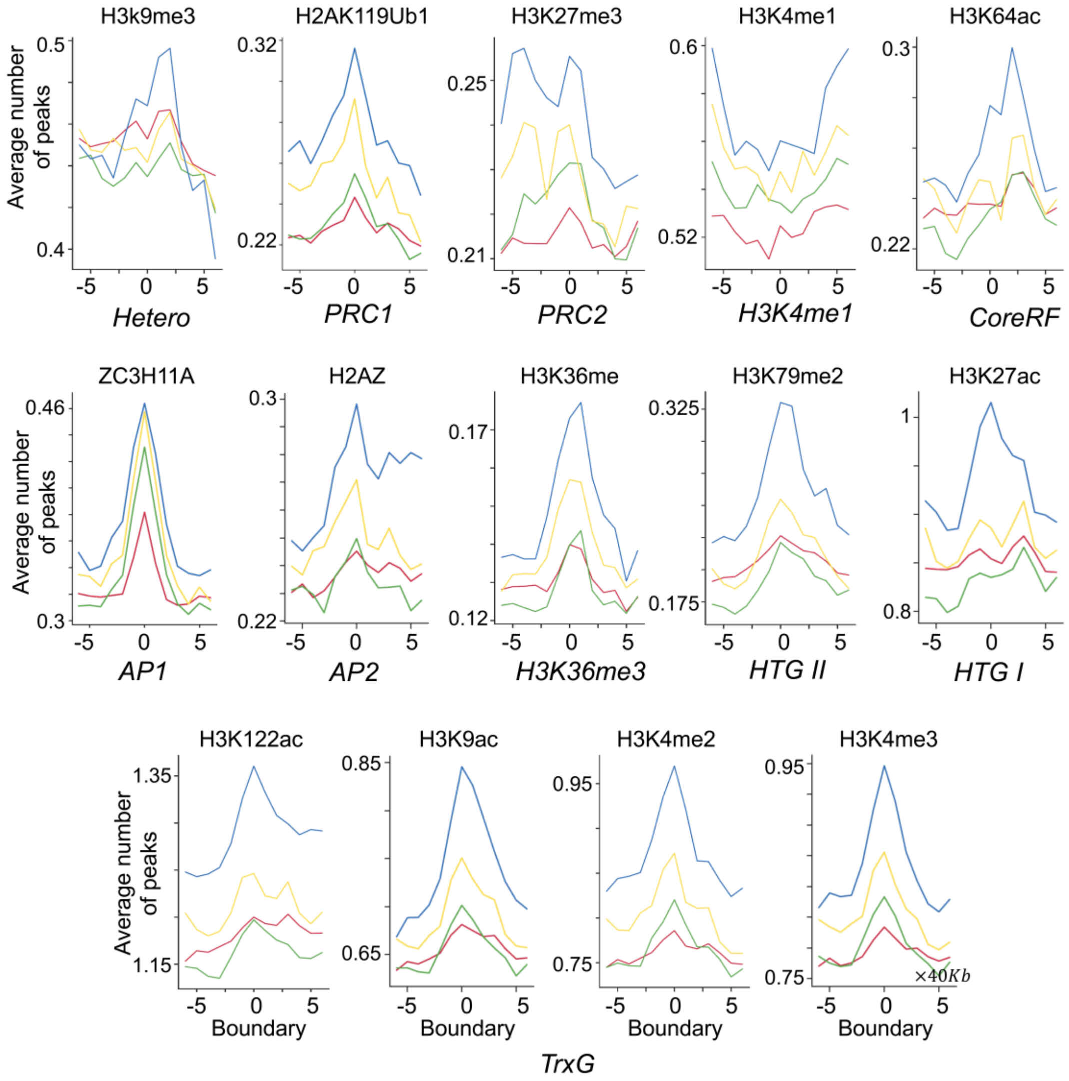


**Figs. S6-S8.** The average number of peaks for other regulatory factors across domain boundaries of all single cells in the different genomic scales. These factors include different classes of regulatory factors annotated in Fig. 4b.

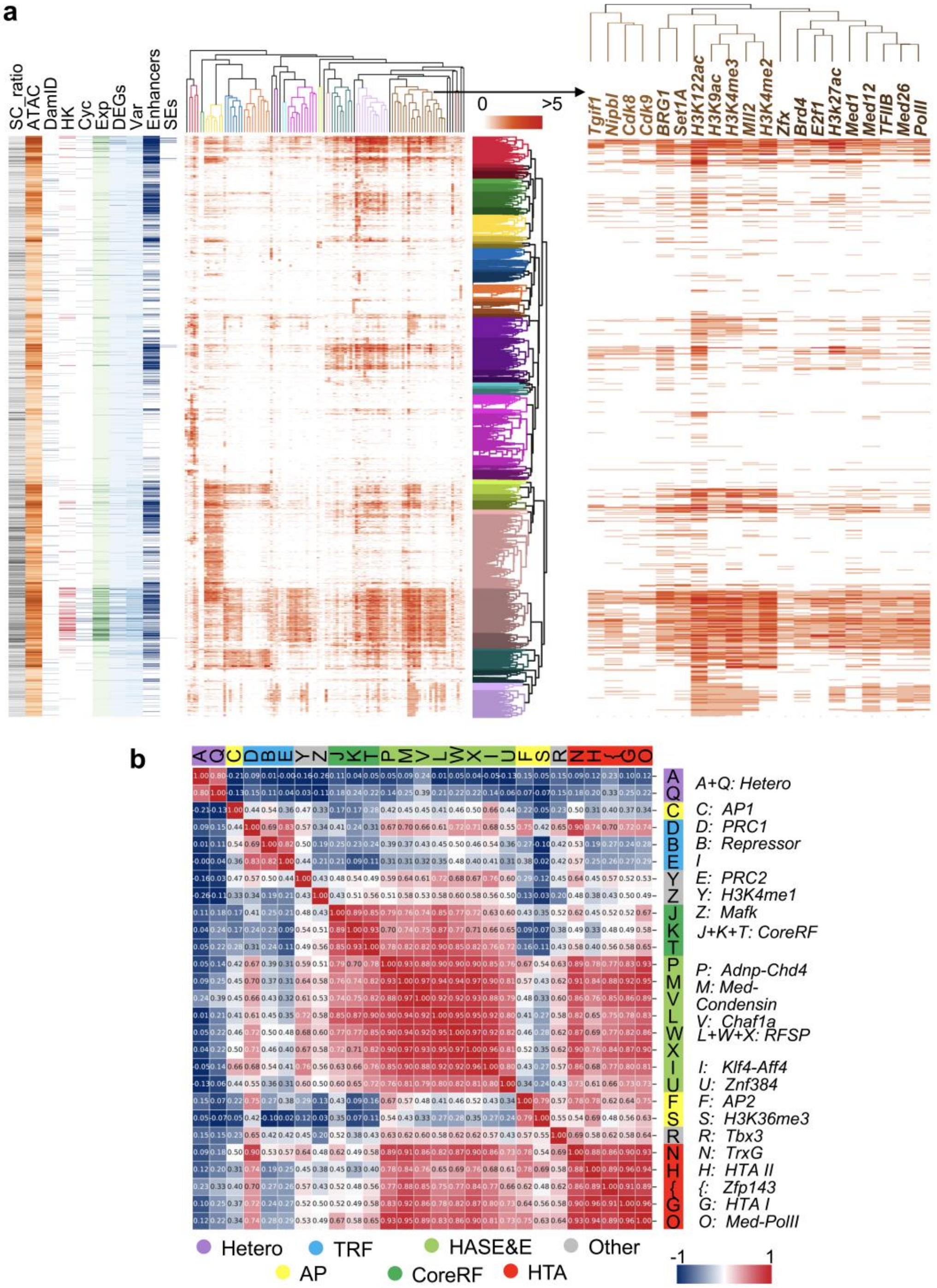


**Fig. S9. a.** The cluster of chromatin positions and regulatory factors. The annotation of chromatin positions by different elements or factors, including SC_ratio (the ratio of single cells forming boundaries), ATAC (ATAC-seq peaks), DamID (DamID-seq), HK (Housekeeping genes), Cyc (mark genes for cell cycle), Exp (gene expression value), DE (differential expression genes), Variable (variable scores for genes), E (enhancers), and SEs (super-enhancers) shown on the left. Hierarchical clustering showing a further division of a certain class of regulatory factors in the right, which is shown in M-O classes of Figure 4a. **b.** The hierarchical clustering showing correlation (left) and annotation (right) of regulatory factor classes.

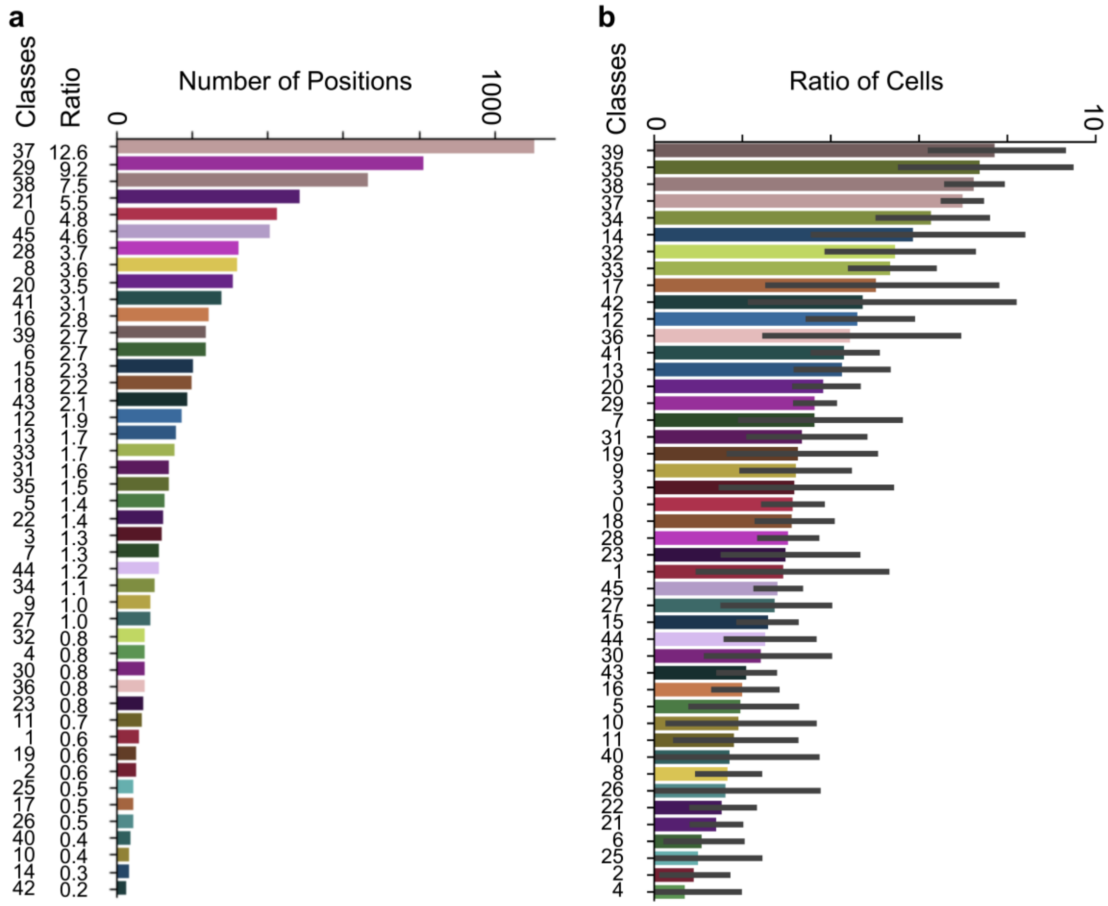


**Fig. S10. a.** The number and ratio of different chromatin position classes. The classes with a ratio>1% were selected for downstream analysis. **b.** The ratio distribution of single cells forming domain boundaries in different chromatin position classes.

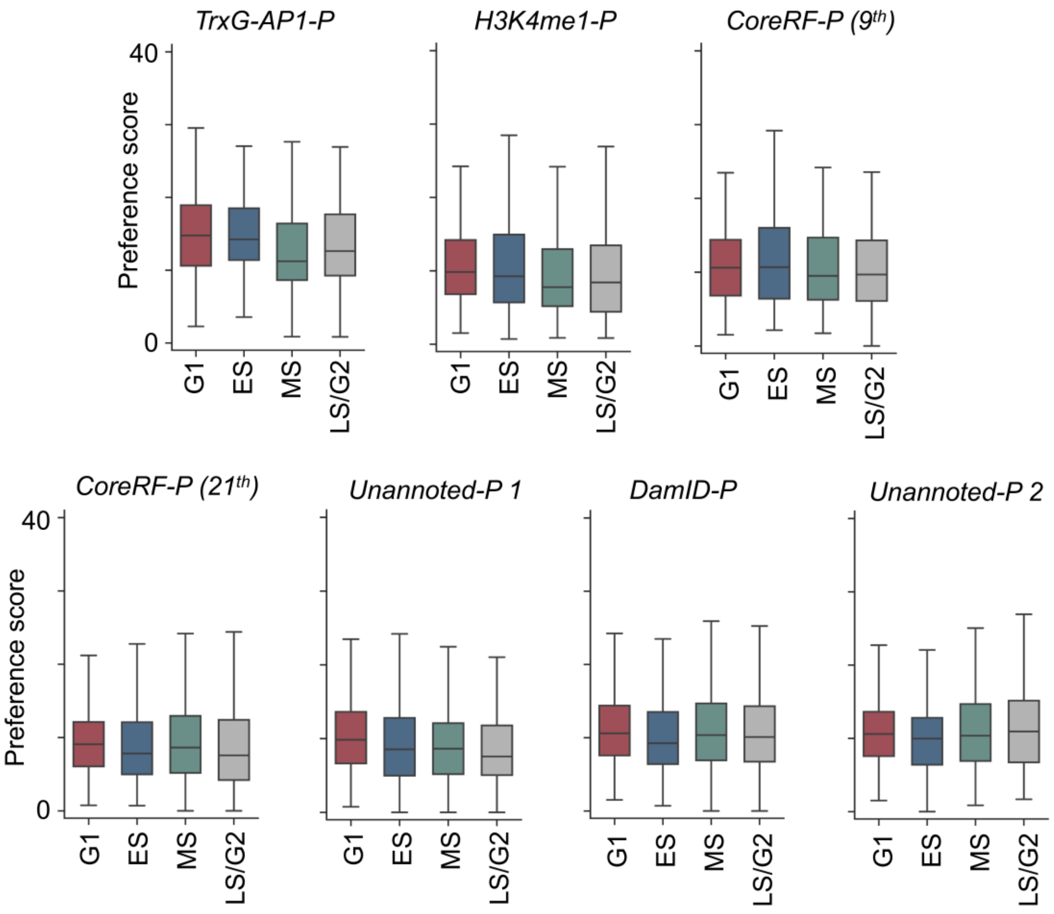


**Fig. S11.** Preference scores of different chromatin landscapes categories across different cell states.

**Supplementary Table Legends**

**Supplementary Table S1.** The source of regulatory factors.
**Supplementary Table S2.** The annotation support of regulatory factor classes.
